# Towards a Kingdom of Reproductive Life – the Core Sperm Proteome

**DOI:** 10.1101/2025.03.06.641666

**Authors:** Taylor Pini, Brett Nixon, Timothy L. Karr, Raffaele Teperino, Adrián Sanz-Moreno, Patricia da Silva-Buttkus, Frank Tüttelmann, Sabine Kliesch, Valérie Gailus-Durner, Helmut Fuchs, Susan Marschall, Martin Hrabě de Angelis, David A. Skerrett-Byrne

**Affiliations:** School of Veterinary Science, The University of Queensland, Gatton, QLD, Australia; Priority Research Centre for Reproductive Science, School of Environmental and Life Sciences, College of Engineering, Science and Environment, The University of Newcastle, Callaghan, NSW, Australia; Hunter Medical Research Institute, Infertility and Reproduction Research Program, New Lambton Heights, NSW, Australia; Biosciences Mass Spectrometry Core Research Facility, Knowledge Enterprise, Arizona State University, USA; ASU-Banner Neurodegenerative Disease Research Center, The Biodesign Institute, Arizona State University, USA; Institute of Experimental Genetics, Helmholtz Zentrum München, German Research Center for Environmental Health, Neuherberg, Germany; German Center for Diabetes Research (DZD) Neuherberg, Germany; Institute of Experimental Genetics, German Mouse Clinic, Helmholtz Zentrum München, German Research Center for Environmental Health, Neuherberg, Germany; Institute of Reproductive Genetics, Centre of Medical Genetics, University and University Hospital of Münster, Germany; Department of Clinical and Surgical Andrology, Centre of Reproductive Medicine and Andrology, University and University Hospital of Münster, Germany; Chair of Experimental Genetics, TUM School of Life Sciences, Technische Universität MuOnchen, Freising, Germany; School of Biomedical Sciences and Pharmacy, College of Health, Medicine and Wellbeing, The University of Newcastle, Callaghan, NSW, Australia

**Keywords:** Sperm, sperm proteome, fertility, data reanalysis, Pebp4, Echs1, Etfb, Ndufa10, Aldh7a1, proteomics, bioinformatics

## Abstract

Reproductive biology is often considered in three siloed research areas; humans, agriculture and wildlife. Yet, each demand solutions for treatment of subfertility, fertility biomarkers, development of assisted reproductive technologies and effective contraception. To efficiently develop solutions applicable to all species, we must improve our understanding of the common biology underpinning reproductive processes. Accordingly, we integrate proteomic data from 29 publicly available datasets (>2 TB of data) to characterize mature sperm proteomes spanning 12 vertebrate species, identifying 13,853 proteins. Although human and mouse have relatively well-annotated sperm proteomes, many non-model species rely heavily on predicted or homology-inferred identifications. Despite variation in proteome size, composition and reproductive strategies, comparative analyses revealed that vertebrates share a fundamental molecular framework essential for sperm function. A core set of 45 species-level and 135 order-level conserved proteins mapped to critical processes, including energy generation, acrosome function, as well as novel signalling pathways (BAG2 and FAT10). Knockout mouse models further validate the significance of these conserved proteins, demonstrating that their disruption impairs sperm motility and fertilization capacity. Moreover, we discovered loss-of-function variants of two additional core sperm proteins in clinical samples, linking them to severe sperm defects. Intriguingly, *in-silico* analysis reveals function-driven, context-dependent diversity surpassing evolutionary patterns. Collectively, these results highlight the value of integrating publicly available datasets and underscore the need for improved genome/proteome annotation in non-model species in mammals. This work provides a foundation for developing cross-species strategies to enhance fertility treatments, assisted reproductive technologies, and conservation efforts. All data is available via ShinySpermKingdom (https://reproproteomics.shinyapps.io/ShinySpermKingdom/).

**IN BRIEF:** Sperm function is essential for fertility across humans, agriculture, and wildlife, yet comparative studies remain limited. This study integrates multi-species proteomic data to identify a core sperm proteome, uncovering conserved molecular pathways and validating novel sperm proteins critical for motility and fertilization.

## INTRODUCTION

Reproductive biology is often considered in three siloed areas: humans, domesticated animals and wildlife. Despite their differences, there are several common needs across these species; efficient production, treatment of subfertility and infertility, development of assisted reproductive technologies (ARTs) and effective contraception (Comizzoli and Holt 2019, Duffy, et al. 2020). To effectively meet these needs, research should focus on developing solutions that are applicable across species where possible. However, to achieve this goal, we need a better understanding of the common reproductive biology across species.

Reproductive physiology is incredibly diverse across the animal kingdom, with many strategies unique to particular evolutionary lineages. In the context of mating, examples of this diversity include the location of semen deposition (Suarez and Pacey 2006), the ability to store spermatozoa in the female tract for extended periods (Holt and Lloyd 2010) and variable sperm morphologies (Fitzpatrick, et al. 2022). Even between species of the same class, some of these traits appear to differ significantly. There is potential to both take advantage of common pathways and potentially exploit strategies that are unique to some species.

This area of study has significant potential benefits on the male side of the reproductive equation. For example, high quality transcriptomic and proteomic studies on the formation of biological sperm storage may highlight pathways that could be targeted to prolong the *in vitro* shelf life of spermatozoa from a variety of species. As an example, koala sperm display an exceptional longevity of up to 42 days post-ejaculation during *ex vivo* storage (Johnston, et al. 2000, Johnston, et al. 2012, Skerrett-Byrne, et al. 2021a). Understanding the biochemical principles of this longevity would yield potential benefits as diverse as human ARTs and addressing the logistical issues of extending the shelf life of fresh semen in the beef and dairy industries (Murphy, et al. 2017). Alternatively, similar studies of spermatozoa and seminal plasma from species with high sperm competition could identify proteins and non-coding RNA species that may be exploited to treat subfertility and improve ARTs. To make such advances, further basic discovery research is required.

With decreasing costs and increasing sensitivity, proteomic profiling of the male gamete has become widespread (Mohanty, et al. 2015). While sperm proteomes have been published for a variety of species, we are yet to capitalize on the suggestion of Oliva, Martínez-Heredia, and Estanyol (Oliva, et al. 2008), to identify conserved proteins from among the wealth of proteomic data that has been collected. As yet there has been limited exploration of cross species analyses; with a single study having compared the sperm proteomes of rodents and ungulates (Bayram, et al. 2016) and another comparing three closely related mouse species with differences in sperm competition (Vicens, et al. 2017). The results of these studies suggest that sperm proteins fall into two categories; (i) highly conserved “core” proteins and (ii) rapidly evolving proteins that are unique to species or taxonomic groups. Importantly, many of the core proteins had important biological roles (e.g. spermatogenesis, capacitation (Bayram, et al. 2016)), suggesting they may provide key avenues for developing cross species reproductive solutions.

A comprehensive cross species sperm proteomic analysis would provide the highest quality data from which to build further research. However, the physical collection of gametes from a large assortment of species involves many difficulties, not the least of which are the considerable expense and logistical management of samples. Thus, as a precursor to further experimental studies, we herein present an *in-silico* analysis of publicly available proteomic data from 12 species, representing the most comprehensive cross species sperm proteome published to date. Using this information, we establish an up-to-date core sperm proteome, highlighting candidate pathways that are highly conserved across species. In addition, we compare proteomes across species based on biological contexts to highlight potentially important pathways for further study.

## MATERIALS AND METHODS

### Chemicals and reagents

Unless otherwise specified, all reagents were purchased from Merck.

### Proteomic data sourcing

Publicly available proteomic data was sourced from the ProteomeXchange repository (www.proteomexchange.org) (Deutsch, et al. 2023), drawing data from a range of partner repositories including PRIDE (Perez-Riverol, et al. 2022), iProX (Chen, et al. 2022), and MassIVE (Choi, et al. 2020). The search term ‘*sperm*’ was initially used to obtain all proteomic datasets containing this keyword. Next, datasets were refined by phylum, with only species in the Chordata phylum retained. In the remaining datasets, any without the RAW mass spectrometry output files (e.g. .WIFF, .RAW) available were excluded. From this filtered list, the remaining datasets were refined based on the following criteria; use of bottom-up proteomics employing data dependent acquisition on fresh, mature cauda epididymal or ejaculated spermatozoa from wildtype animals. Studies were not excluded based on sample enrichment techniques (e.g. density gradient centrifugation, isolation of plasma membrane proteins). Beyond these criteria, studies were excluded if the experimental treatment of each raw MS output file could not be determined or available RAW files deposited were not of sufficient detail or file type for reanalysis. In the first case, clarification was sought from the publishing authors to establish file identities prior to exclusion. The application of these criteria resulted in 29 datasets (Table 1) and over 2TB of RAW data.

**Table 1.** Final proteomic datasets included in this study.

| **Study** | **Dataset identifier** | **PMID** | **Species** |
| --- | --- | --- | --- |
| (Byrne, et al. 2012) | PXD000007 | 23081703 | *Bos indicus* |
| (Ramesha, et al. 2020) | PXD014172 | 32508098 | *Bos indicus* |
| (Kasvandik, et al. 2015) | PXD001096 | 25603787 | *Bos taurus* |
| (Shen, et al. 2021) | PXD019435 | 34009990 | *Bos taurus* |
| (Fu, et al. 2019) | PXD003859 | 31146187 | *Bubalus bubalis* |
| (Batra, et al. 2021) | PXD022114 | 34174811 | *Bubalus bubalis* |
| (Nixon, et al. 2019) | MSV000082258 | 30072580 | *Crocodylus porosus* |
| (Labas, et al. 2015) | PXD001254 | 25086240 | *Gallus gallus* |
| (Vitorino Carvalho, et al. 2021) | PXD022322 | 33898456 | *Gallus gallus* |
| (Vandenbrouck, et al. 2016) | PXD003947 | 27444420 | *Homo sapiens* |
| (Schiza, et al. 2018) | PXD007515 | 30097533 | *Homo sapiens* |
| (Urizar-Arenaza, et al. 2019) | PXD011290 | 30622161 | *Homo sapiens* |
| (Pini, et al. 2020) | PXD014849 | 32026202 | *Homo sapiens* |
| (Castillo, et al. 2019) | PXD014871 | 31824947 | *Homo sapiens* |
| (Guyonnet, et al. 2012) | PXD000575 | 22707618 | *Mus musculus* |
| (Guyonnet, et al. 2014) | PXD000592 | 24797071 | *Mus musculus* |
| (Castaneda, et al. 2017) | PXD005343 | 28630322 | *Mus musculus* |
| (Liu, et al. 2019) | PXD013092 | 30867229 | *Mus musculus* |
| (Xu, et al. 2020) | PXD016928 | 31969357 | *Mus musculus* |
| (Skerrett-Byrne, et al. 2022) | PXD028834 | 36384108 | *Mus musculus* |
| (Casares-Crespo, et al. 2019) | PXD007989 | 30753958 | *Oryctolagus cuniculus* |
| (Juárez, et al. 2020) | PXD015510 | N/A | *Oryctolagus cuniculus* |
| (Leahy, et al. 2020) | PXD017537 | 32383290 | *Ovis aries* |
| (Skerrett-Byrne, et al. 2021a) | PXD024250 | 34411425 | *Phascolarctos cinereus* |
| (Xu, et al. 2021) | PXD025607 | 34336820 | *Sus scrofa* |
| (Pérez-Patiño, et al. 2019) | PXD010062 | 30257877 | *Sus scrofa* |
| (Zhang, et al. 2022) | PXD030020 | 35252197 | *Sus scrofa* |
| (Fuentes-Albero, et al. 2021) | PXD024588 | 34336830 | *Tursiops truncatus* |
| (Bayram, et al. 2016) | PXD003164 | 26768581 | *Mus musculus, Bos taurus, Sus scrofa, Rattus norvegicus* |

### International Mouse Phenotyping Consortium mouse models, histology, and data collection

The International Mouse Phenotyping Consortium (IMPC) database (Dickinson, et al. 2016, Groza, et al. 2022) was mined for genetic knockout mice overlapping with those identified as the core sperm proteome (Fig. 3A), and with special access granted, we crossed referenced with the European Mouse Mutant Archive (EMMA) (Hagn, et al. 2007) to restrict to those gene KOs with available *in vitro* fertilization (IVF) and sperm data. The mouse models were generated using the IMPC targeting strategy with CRISPR/Cas technology at Helmholtz Munich (https://www.mousephenotype.org/understand/the-data/allele-design/). After genotyping, heterozygous × heterozygous matings were set up to generate sufficient mutant mice with littermate ^+/+^ controls for phenotyping analysis at the German Mouse Clinic as described, (Fuchs, et al. 2018) and in agreement with the standardized phenotyping pipeline of the IMPC including histopathological analysis (n=2) (https://www.mousephenotype.org/impress/PipelineInfo?id=14) for all lines except for *Aldh7a1* (n=5). We obtained data pertinent to *Aldh7a1* (Aldehyde dehydrogenase 7 family member A1), *Echs1* (Enoyl-CoA hydratase, short chain 1), *Etfb (*Electron transfer flavoprotein subunit beta), *Ndufa10* (NADH:ubiquinone oxidoreductase subunit A10), *Pebp4* (Phosphatidylethanolamine binding protein 4), with a wildtype reference control provided by EMMA. For further information on protocols used by EMMA for sperm collection, analysis and IVF, please see their publicly available resources and videos (https://www.infrafrontier.eu/emma/cryopreservation-protocols/). Briefly, histopathological analyses of formalin-fixed, paraffin-embedded and haematoxylin & eosin (H&E)-stained sections (3 μm-thick) of testis, epididymis, prostate and seminal vesicles from control and mutant mice were performed blind by two pathologists. When applicable, the number of multinucleated giant cells (MGCs) in the seminiferous tubules was counted and expressed per unit area of testis.

### Proteome Discoverer processing

Consistent with previous studies (Martin, et al. 2022, Murray, et al. 2021, Skerrett-Byrne, et al. 2022, Skerrett-Byrne, et al. 2021a, Skerrett-Byrne, et al. 2021b, Skerrett-Byrne, et al. 2021c, Smyth, et al. 2022, Staudt, et al. 2022, Trigg, et al. 2021), database searching of each study’s RAW files was performed using Proteome Discoverer 2.5 (Thermo Fisher Scientific). SEQUEST HT was used to search against the appropriate UniProt database (Table S1, each downloaded 18^th^ June 2022, including reviewed and unreviewed proteins). Highly stringent database searching criteria were utilized, including up to two missed cleavages, a precursor mass tolerance set to 10 ppm and fragment mass tolerance of 0.02 Da. Trypsin was designated as the digestion enzyme. Cysteine carbamidomethylation was set as a fixed modification while acetylation (K, N-terminus), phosphorylation (S,T,Y) and oxidation (M) were designated as dynamic modifications. Interrogation of the corresponding reversed database was also performed to evaluate the false discovery rate (FDR) of peptide identification using Percolator on the basis of *q*-values, which were estimated from the target-decoy search approach. To filter out target peptide spectrum matches over the decoy-peptide spectrum matches, a fixed FDR of 1% was set at the peptide level. The resultant protein list was exported from Proteome Discoverer 2.5 as an Excel file and further refined to include only those with a protein identification (FDR ≤ 0.01) with at least one or more unique peptides.

### Phylogenetic trees and UniProt mapping

NCBI taxonomy numbers were submitted to phylot (v2) to generate a phylogenetic tree, visualized and exported from iTOL (Interactive Tree Of Life) (Letunic and Bork 2007, 2011). Utilising UniProt (https://www.uniprot.org/), each of the sperm proteomes were mapped to the UniProt Knowledge Base to ascertain the level of evidence of each protein (Skerrett-Byrne, et al. 2022, Skerrett-Byrne, et al. 2021a, Skerrett-Byrne, et al. 2021b, Skerrett-Byrne, et al. 2021c, Smyth, et al. 2022). UniProt protein evidence is a measure of the current, manually curated, type of evidence that supports the existence of that protein; experimental evidence 1) at protein level; 2) at transcript level; 3) protein inferred from homology; 4) protein predicted.

### Humanization with OmicsBox

To conduct a comparative analysis between all species, a minimum cut-off of at least 500 proteins was applied before proceeding to humanization to maximize the comparisons possible. Data from each of the remaining eight species were uploaded to UniProt to generate a FASTA file for humanization using a custom workflow on the OmicsBox software (version 2.2.4, BioBam Bioinformatics, Valencia, Spain) (https://www.biobam.com/omicsbox). This workflow includes a cloud based DIAMOND BLAST protein search against the human proteome (Buchfink, et al. 2021, Götz, et al. 2008, Skerrett-Byrne, et al. 2021a, Zhang, et al. 2022), with the output restricted to an e-value cut-off of 4.07E^-10^ to ensure accurate homologues were obtained (97.5% average conversion).

### Identifying conserved and species-specific proteins

Conserved proteins were identified at both species level (i.e., proteins present in all species used for further analysis) and order level (i.e., proteins present in at least one species of all taxonomic orders used for further analysis). Humanized identifications (IDs) were employed for this analysis and lists were compared using jvenn (Bardou, et al. 2014) and DeepVenn (Hulsen 2022) to identify conserved proteins. The analysis at order level included the taxonomic orders Primates (*H. sapiens*), Rodentia (*M. musculus*), Artiodactyla (*B. taurus, O. aries, S. scrofa, T. truncates*), Lagomorpha (*O. cuniculus*), Diprotodontia (*P. cinereus*) and Crocodilia (*C. porosus*).

### Comparing the sperm proteome based on biological contexts

Groups of species were compared based on several biological ‘contexts’, including location of testes (internal vs external), history of selective breeding (yes vs no) and sperm metabolism preference (glycolysis preference vs no preference). Humanized IDs were used for this analysis and species classified into each group are listed in Supplementary Table S17. To account for the stronger influence of human and mouse proteomes due to their extensive inventories, proteins were not included in the analysis if they were only identified in mouse or human spermatozoa. Lists were compared using jvenn (Bardou, et al. 2014) and DeepVenn (Hulsen 2022) to identify conserved and unique proteins.

### Bioinformatic analyses of proteomic data

Bioinformatic analyses employed humanized IDs for analysis. High granularity pathway analysis was performed using the Ingenuity Pathway Analysis software package (IPA; Qiagen, Hilden, Germany) as previously described (Martin, et al. 2022, Murray, et al. 2021, Skerrett-Byrne, et al. 2022, Skerrett-Byrne, et al. 2021a, Skerrett-Byrne, et al. 2021b, Skerrett-Byrne, et al. 2021c, Smyth, et al. 2022, Staudt, et al. 2022, Trigg, et al. 2021, Zhang, et al. 2022). Each humanized proteomic list was analysed on the basis of predicted protein subcellular location and classification (other excluded), in addition to canonical pathways and disease and functions, using the IPA *p*-value enrichment score (a strict cut-off of *p*-value ≤ 0.05) (Krämer, et al. 2013). The Database for Annotation, Visualization and Integrated Discovery (DAVID, www.david.ncifcrf.gov, v 2021, (Huang da, et al. 2009, Sherman, et al. 2022)) functional annotation clustering tool was used to identify enriched clusters based on gene ontology terms, protein-protein interactions, protein domains, pathways and literature. All searches were performed with default thresholds for similarity, classification and enrichment, using *Homo sapiens* as the background gene list. Clusters were classified as significantly enriched based on Benjamini adjusted *p*-values ≤ 0.05. Visual protein-protein interaction networks were generated using STRING (www.string-db.org, v 11.5). The humanized proteome was interrogated using UniProt to assess subcellular locations relevant to sperm cells, the following GO terms were used: acrosomal vesicle (GO:0001669), perinuclear theca (GO:0033011), nucleus (GO:0005634), mitochondrion (GO:0005739), axoneme (GO:0005930), cytoskeletal calyx (GO:0033150), sperm midpiece (GO:0097225), head-tail coupling apparatus (GO:0120212), principal piece (GO:0097228), end piece (GO:0097229), annulus (GO:0097227) and flagellum (GO:0036126).

### Clustering and network visualization

The refined pathways output from IPA were loaded into Perseus (version 1.6.10.43) (Tyanova, et al. 2016), to carry out unbiased hierarchical clustering across the species. Protein networks of the core sperm proteomes at the species (45 proteins) and order (135 proteins) taxonomic levels were investigated using STRING (version 12.0)(Szklarczyk, et al. 2021) and then visualized and modified using Cytoscape (version 3.8.2). (Shannon, et al. 2003) Basic data handling, if not otherwise stated, was conducted using Microsoft Excel 365 (Version 2211, Microsoft Corporation, Redmond, WA) and GraphPad Prism version 10.4.1 (GraphPad Software; San Diego, CA).

### Shiny Application development

In accordance with Shiny blueprint outlined by ShinySperm(Skerrett-Byrne, et al. 2024), a Shiny Application was deployed to support the accessibility and interpretability of these datasets within, allowing for effective data-driven insights by the field. The full coding script supporting ShinySpermKingdom (https://reproproteomics.shinyapps.io/ShinySpermKingdom/), can be downloaded from GitHub – https://github.com/DavidSBEire/ShinySpermKingdom. In brief, the ShinySpermKingdom application was built using the shiny package (version 1.9.1) on RStudio (version 2024.04.1+748), with base *R* (version 4.3.3, 2024-02-29). Supporting the functionality and aesthetics of this application are several packages, including: DT, eulerr, ggplot2, openxlsx, plotly, readxl, reshape2, RColorBrewer, and shinydashboard.

### Male Reproductive Genomics (MERGE) cohort

The MERGE cohort currently comprises exome/genome data of almost 3,000 men of whom most attended the Centre of Reproductive Medicine and Andrology (CeRA), Münster, for couple infertility. MERGE is continuously growing and for the current study, 2,882 datasets of men with quantitative and/or qualitative sperm defects were queried for the 135 genes encoding the identified conserved sperm proteins. Specifically, 2,327 men had very few or no sperm in the ejaculate (crypto-or azoospermia, HP:0030974/HP:0000027), 437 had various grades of reduced sperm counts (oligozoospermia, HP:0000798) often combined with reduced/impaired sperm motility and/or morphology (astheno-/teratozoospermia, HP:00122077/HP:0012864), and 118 had normal sperm counts but motility and/or morphology defects. The most recent description of MERGE including the details of sequencing are available in (Stallmeyer, et al. 2024). Only well-covered (>20x), rare (minor allele frequency [MAF] <0.01 in gnomAD 2.1.1), coding, homo-or hemizygous, loss-of-function variants (LoF: stop gained, frameshift, splice acceptor/donor) were prioritised.

## RESULTS

### Establishment of multispecies mature spermatozoa proteomes

A comprehensive search of the ProteomeXchange repository using the keyword ‘*sperm*’ yielded a total of 146 datasets (Fig. 1A). After excluding species from outside the Chordata phylum, a total of 90 datasets remained. Of these, 48 studies were excluded based on our predefined inclusion criteria, such as only studies on functional mature sperm cells (See STAR methods). A further 13 studies were excluded based on insufficient information available for reanalysis. Datasets at each successive level of exclusion are listed in Table S1.

**Figure 1:**
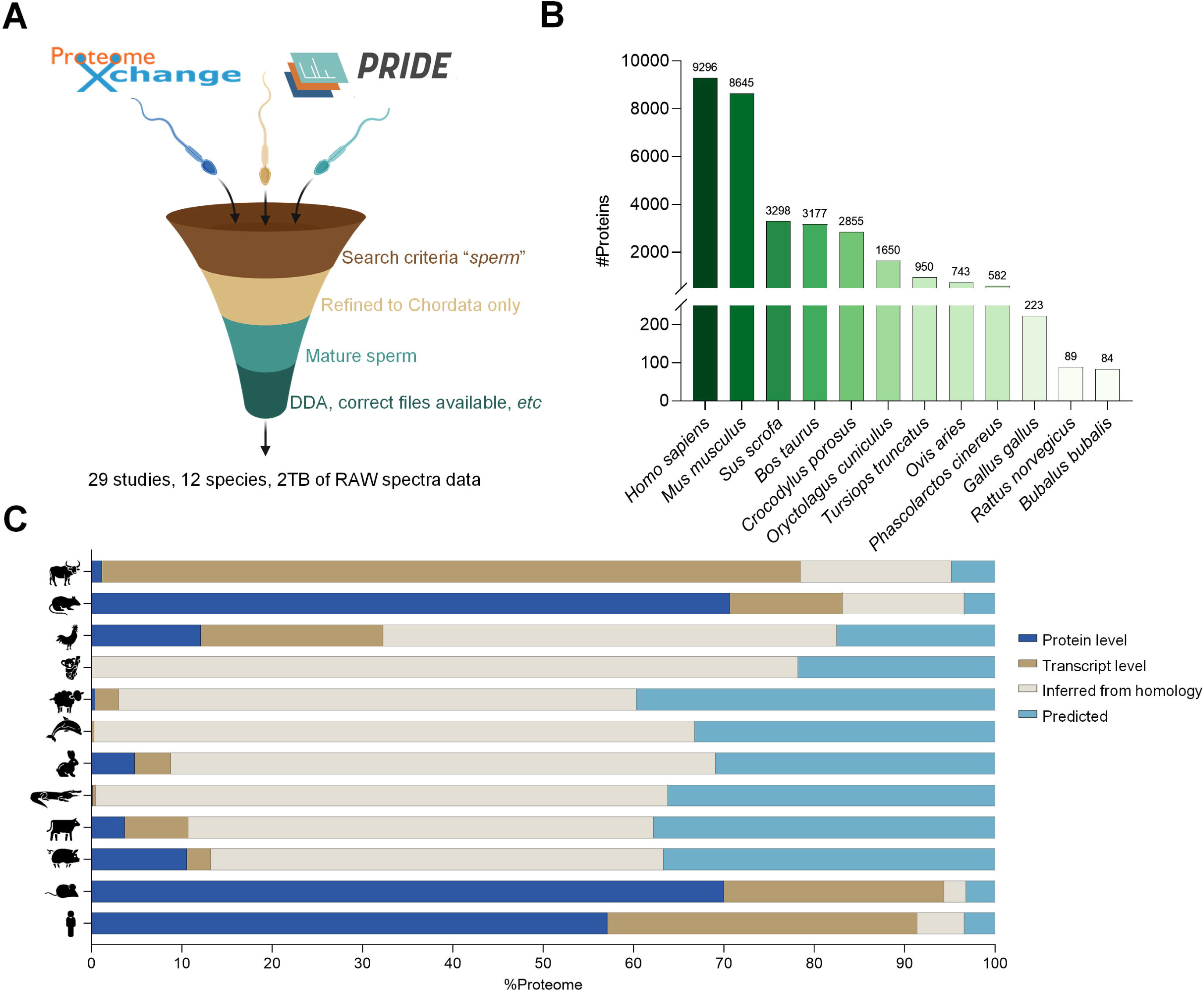
Characterization of multispecies sperm proteomes. (A) Across 12 different species, over 2 TB of RAW spectral data was sourced from public repositories and processed using Proteome Discoverer 2.5, utilising highly stringent criteria. (B) The number of proteins identified in each of the 12 species and (C) the proportion their respective level of evidence for protein evidence as curated by to the UniProt Knowledge Base; 1) at protein level; 2) at transcript level; 3) protein inferred from homology; 4) protein predicted.

The final cohort comprised 29 datasets, representing 12 species: *Homo sapiens* (Castillo, et al. 2019, Pini, et al. 2020, Schiza, et al. 2018, Urizar-Arenaza, et al. 2019, Vandenbrouck, et al. 2016), *Mus musculus* (Bayram, et al. 2016, Castaneda, et al. 2017, Guyonnet, et al. 2014, Guyonnet, et al. 2012, Liu, et al. 2019, Skerrett-Byrne, et al. 2022, Xu, et al. 2020), *Sus scrofa* (Bayram, et al. 2016, Pérez-Patiño, et al. 2019, Xu, et al. 2021, Zhang, et al. 2022), *Bos taurus* (Bayram, et al. 2016, Byrne, et al. 2012, Kasvandik, et al. 2015, Ramesha, et al. 2020, Shen, et al. 2021), *Crocodylus porosus* (Nixon, et al. 2019), *Oryctolagus cuniculus* (Casares-Crespo, et al. 2019, Juárez, et al. 2020), *Tursiops truncatus* (Fuentes-Albero, et al. 2021), *Ovis aries* (Leahy, et al. 2020), *Phascolarctos cinereus* (Skerrett-Byrne, et al. 2021a), *Gallus gallus domesticus* (Labas, et al. 2015, Vitorino Carvalho, et al. 2021), *Rattus norvegicus* (Bayram, et al. 2016) and *Bubalus bubalis* (Batra, et al. 2021, Fu, et al. 2019) (Fig. 1A, Table S1). These datasets were reanalysed using a stringent and uniform pipeline implemented in Proteome Discover, and the resultant protein IDs are provided in Tables S2-13. To enhance accessibility, all data is also available on ShinySpermKingdom, facilitating an interactive experience with these complex datasets (https://reproproteomics.shinyapps.io/ShinySpermKingdom/).

Unsurprisingly, human (9,296 proteins) and mouse (8,645 proteins) exhibited the most comprehensive sperm proteomes (Fig. 1B, Table 2), reflecting their status as well researched species. Returning nearly a third of the larger proteomes was the boar (3,298), closely followed by bull (3,177), crocodile (2,855), and rabbit (1,650). The proteome of each species was first assessed using UniProt to determine their current curated level of protein evidence (Fig. 1C; Tables S2-13). Predictably, the sperm proteomes of well-characterized species like human (91.4%), mouse (94.4%) and rat (83.1%) were all well annotated at protein and transcript level. Interestingly, buffalo sperm harboured 77.3% of its evidence at the transcript level. Within the sperm proteomes of the remaining 8 species, in most cases >90% of protein identifications were only predicted or inferred from homology, indicating that experimental evidence for the existence of most proteins remains poor in non-traditional model species (Fig. 1C). Due to the low number of protein identifications in the chicken (223 IDs), Norwegian rat (89 IDs) and buffalo (84 IDs), these species were excluded from further downstream analyses (Fig. 1B). These exclusions ensured a focus on datasets with sufficient coverage and quality for robust comparative analysis.

**Table 2.** Summary of protein identifications by species.

| Species | #Studies | #Original<br>protein IDs | Humanized<br>IDs | Conversion<br>rate (%) | # Unique to<br>species (%) |
| --- | --- | --- | --- | --- | --- |
| Human (*Homo sapiens*) | 5 | 9296 |  |  | 6093 (66.3) |
| House mouse (*Mus musculus*) | 7 | 8645 | 8424 | 97.4 | 2536 (30.1) |
| Boar (*Sus scrofa*) | 4 | 3298 | 3190 | 96.7 | 111 (3.5) |
| Cattle (*Bos taurus/indicus*) | 5 | 3177 | 3048 | 95.9 | 149 (4.9) |
| Saltwater crocodile (*Crocodylus porosus*) | 1 | 2855 | 2743 | 96.1 | 165 (6.0) |
| European rabbit (*Oryctolagus cuniculus*) | 2 | 1650 | 1633 | 99.0 | 59 (3.6) |
| Common bottlenose dolphin (*Tursiops truncates*) | 1 | 950 | 941 | 99.1 | 15 (1.6) |
| Sheep (*Ovis aries*) | 1 | 743 | 726 | 97.7 | 40 (5.5) |
| Koala (*Phascolarctos cinereus*) | 1 | 582 | 573 | 98.5 | 26 (4.5) |
| Chicken (*Gallus gallus*) | 2 | 223 |  |  |  |
| Brown rat (*Rattus norvengicus*) | 1 | 89 |  |  |  |
| Domestic water buffalo (*Bubalus bubalis*) | 2 | 84 |  |  |  |

### Humanized sperm proteomes redefine evolutionary links

To advance the understanding of the remaining 9 species, each sperm proteome was converted to their respective human homologues to allow utilization of human focused bioinformatic tools, facilitating a standardized cross species analysis. Humanization was achieved using the OmicsBox software as previously described.(Skerrett-Byrne, et al. 2021a) Conversion rates to humanized IDs were exceptionally high, ranging from 95.9% – 99.1%, producing a total of 13,853 proteins (Fig. 2A, Table 2, Table S14). These newly generated sperm proteomes were subject to analyses using Ingenuity Pathway Analysis (IPA) to provide an overall classification of protein types present. Notably, despite variations in proteome size, proportional compositional analysis of each species revealed broad consistency (Fig. 2B). Enzymes were the dominated category, accounting for ∼72.5% of all sperm proteins, followed by transporters (∼15.5%), transcription (∼5.3%) and translation (∼2.7%) regulators, receptors (∼1.6%), ion channels (∼1.6%), cytokines (∼0.6%) and growth factors (∼0.5%).Whilst there were no overt differences in the proportional composition of protein classification types, the two largest sperm proteomes (i.e., human and mouse), featured proportionally more transcription regulators than that of all other species assessed (i.e., ∼9.3% vs ∼4.1%). This enrichment underscores potential differences in sperm-specific regulatory mechanisms between traditional model organisms and less-studied species. Further interrogation with UniProt subcellular localization and the Human Protein Atlas(Uhlén, et al. 2015) sperm subcellular resource allowed mapping to key sperm locations (Fig. S1): acrosome (146 proteins), perinuclear theca (17 proteins), nucleus (3,879 proteins), calyx (15 proteins), equatorial segment (13 proteins), connecting piece (45 proteins), mid-piece (118 proteins), mitochondria (394 proteins), annulus (16 proteins), flagellum (204 proteins), flagellar centriole (21 proteins), axoneme (176 proteins), principal piece (91 proteins), and end piece (31 proteins) (Table S14).

**Figure 2:**
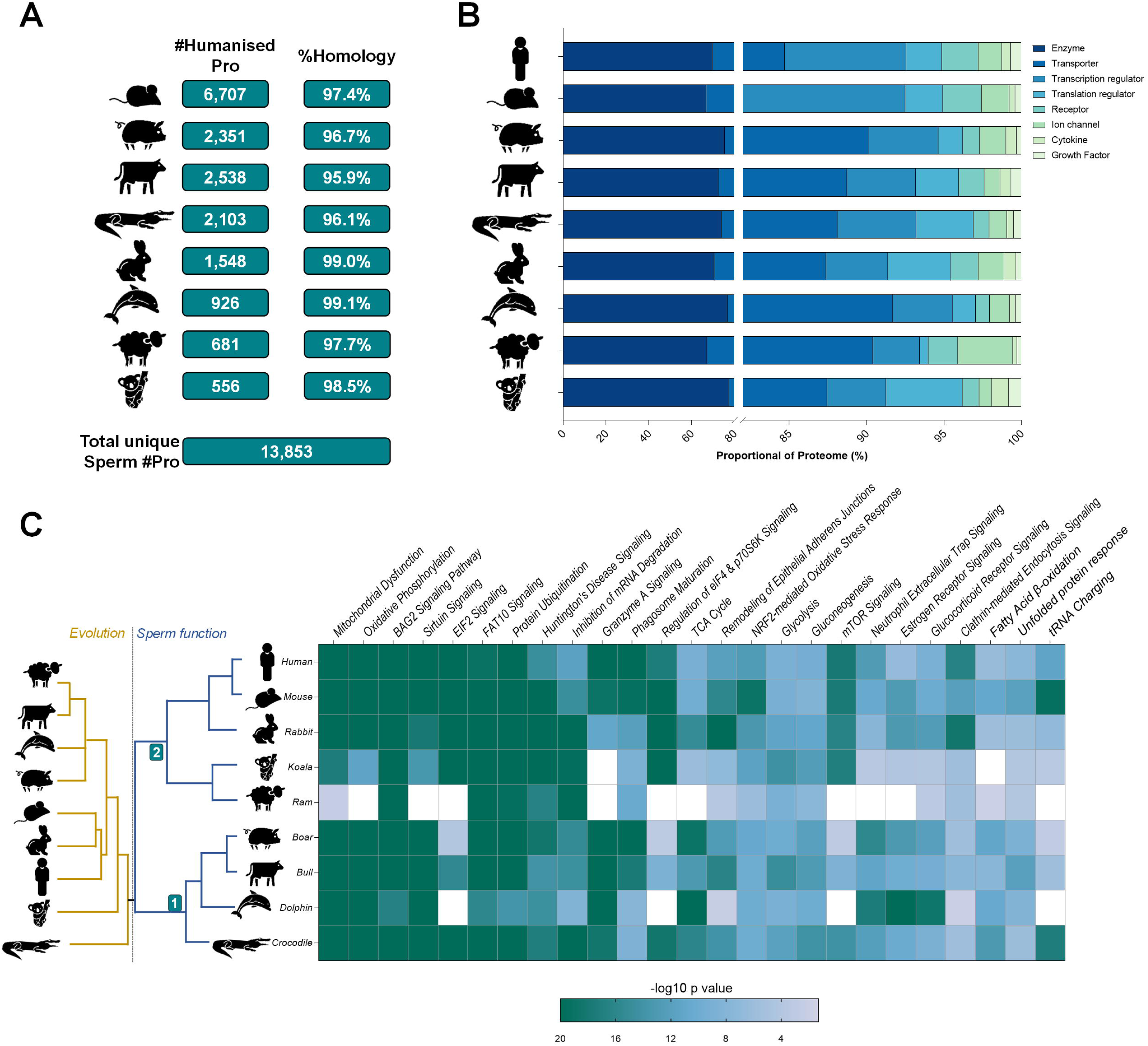
Humanization of sperm proteomes. (A) Summary of the number of proteins which were successfully converted to human homologues (duplicates removed). (B) The percentage of the original species proteome retained. (C) Heatmap depicts the top 25 unique pathways significantly enriched in at least one species (*p*-value ≤ 0.05), white denoting absence of detection. To the left, in gold is a phylogenetic tree depicts the evolutionary distances and relationships between the 9 species (generated with phylot and iTOL). Mirrored to the right in blue is the unbiased hierarchical clustering based upon the full remit of 482 pathways (sperm function).

Seeking to delve further into the functional relationships among these sperm proteomes, the canonical pathways node of IPA was utilized leading to the identification of 482 unique pathways significantly enriched in at least one species *(p*-value ≤ 0.05). Unbiased hierarchical clustering based on the conservation of these pathways revealed mouse spermatozoa to be the most functionally related to that of their human counterparts (Fig. 2C). This finding contrasts that of the genomic lineage tracing, which indicated that among the species assessed, rabbits were the closest evolutionary relative to humans. Notably, this hierarchical clustering approach achieved the division of the assessed species into two broad groupings; 1) crocodile, dolphin, bull and boar; 2) sheep, koala, rabbit, mouse and human. This reorganization suggests that functional relationships based on sperm proteomes may not strictly align with evolutionary distances derived from genomic data. Amongst the most significantly enriched pathways across all species were those related to energy metabolism (“Mitochondrial Dysfunction”, “Oxidative Phosphorylation”, “Glycolysis”, “Gluconeogenesis”, and “Fatty Acid β-oxidation”) and capacitation (“Sirtuin Signalling Pathway”, “Protein Ubiquitination Pathway”).

### Characterization of the core sperm proteome

To ascertain the proteins most fundamental to the functional competency of spermatozoa across the 9 assessed species, a comparison of all protein identifications was conducted. This strategy uncovered a modest 45 and 135 conserved proteins at the taxonomic level of species and orders, respectively, which we hereafter refer to as the core sperm proteome (Fig. 3A, Table S15). The 9 species were collapsed into six taxonomic orders, namely Primate (human; 9,186 proteins), Rodentia (mouse; 6,707 proteins), Artiodactyl (boar, bull, dolphin and sheep; 3,795 proteins), Lagomorpha (rabbit; 1,548 proteins), Diprotodontia (koala; 556 proteins), and Crocodilia (crocodile; 2,103 proteins). Of those proteins that were identified in more than one order, the largest overlaps were between Primate and Rodentia (830), Primate, Rodentia and Artiodactyla (489), and Rodentia and Artiodactyla (457) (Fig. 3A).

**Figure 3:**
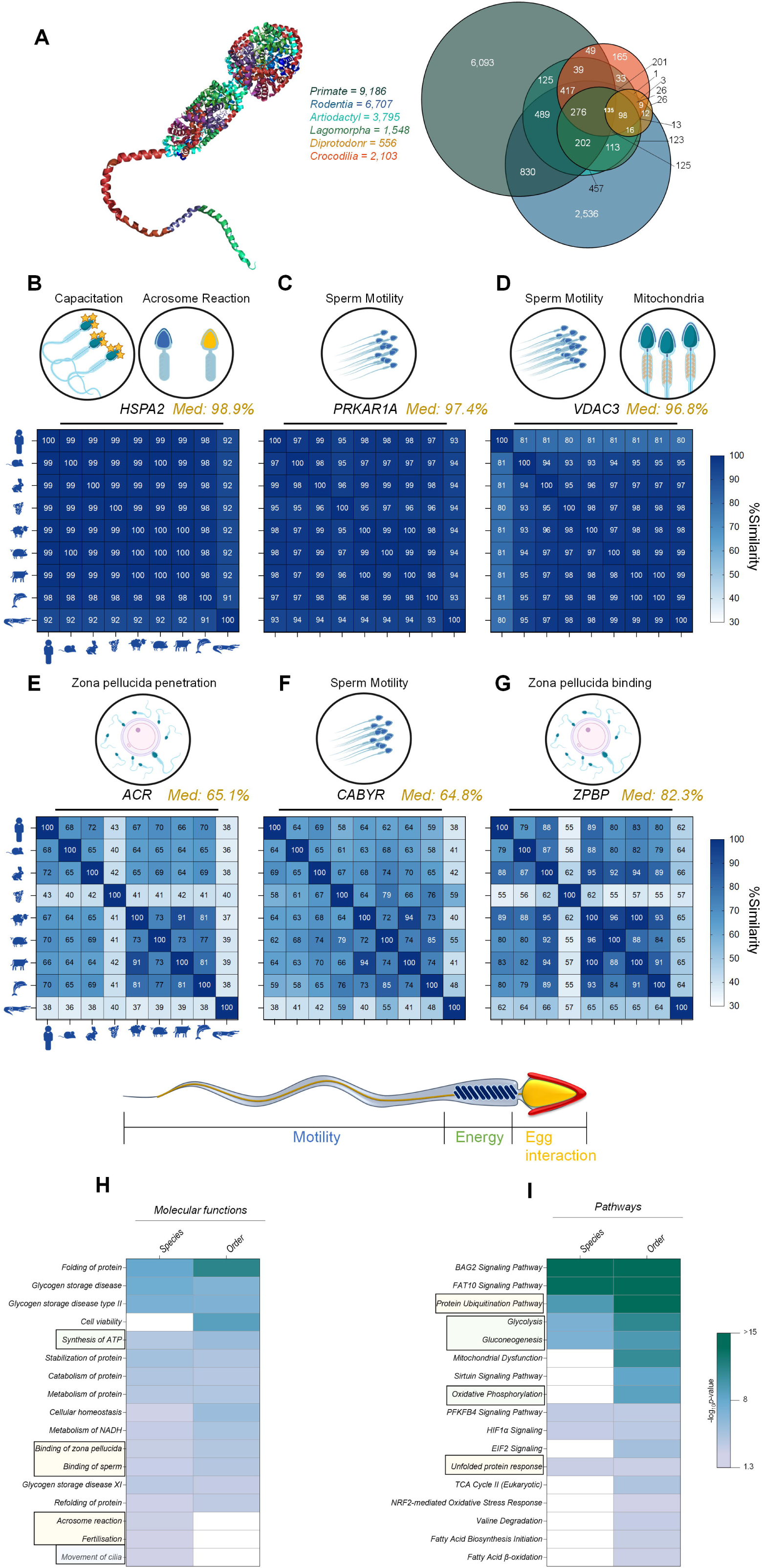
Core sperm proteome. (A) Venn Diagram depicts the overlap of sperm proteomes at the level of taxonomic orders; Primates (H. sapiens), Rodentia (M. musculus), Artiodactyla (B. taurus, O. aries, S. scrofa, T. truncates), Lagomorpha (O. cuniculus), Diprotodontia (P. cinereus) and Crocodilia (C. porosus). UniProt Alignment tool (Clustal Omega) was utilized to interrogate the sequence alignment between each species for (B) Heat shock protein family A member 2 (HSPA2), (C) Protein kinase cAMP-dependent type I regulatory subunit alpha (PRKAR1A), (D) Voltage dependent anion channel 3 (VDAC3), (E) Acrosin (ACR), (F) Calcium binding tyrosine phosphorylation regulated (CABYR), and (G) Zona pellucida binding protein (ZPBP). The median alignment (% Similarity) is denoted in gold next to each protein symbol. The conserved sperm proteomes at the level of species (45 proteins) and order (135 proteins) were subjected to analysis using Ingenuity Pathway Analysis (IPA). Heatmaps depict the comparative analysis of species and orders, with a refined focus on reproductive related (H) molecular functions and (I) pathways. Pathways and function known to important to motility, energy and egg interactions are highlight by blue, green and gold boxes respectively.

Further interrogation of these conserved proteins with the UniProt Alignment tool (Clustal Omega(Sievers, et al. 2011)) revealed many proteins with known roles in sperm motility, mitochondria function, capacitation, and acrosome reaction have high levels of sequence similarity (Fig. 3B–D); heat shock protein family A member 2 (HSPA2; 98.9%), protein kinase cAMP-dependent type I regulatory subunit alpha (PRKAR1A; 97.4%), and voltage dependent anion channel 3 (VDAC3; 96.8%). However, a notable divergence was observed for human VDAC3, which differs by ∼20% compared to the other 8 species. Conversely, proteins critical for sperm motility, zona pellucida binding, and penetration exhibited greater variability in sequence conservation (Fig. 3E–G); acrosin (ACR, 65.1%), calcium binding tyrosine phosphorylation regulated (CABYR; 64.8%), and the zona pellucida binding protein (ZPBP; 82.3%).

Both species and order lists were subjected to analysis with IPA, focusing on molecular functions and pathways. Strong consistency between both groups was observed, with the significant enrichment (*p*-value ≤ 0.05) of key reproductive processes, including synthesis of ATP, movement of cilia, acrosome reaction, binding of sperm and zona pellucida, and fertilization (Fig. 3H, Table S16). Pathway analysis of the core sperm proteomes displayed significant enrichment of pathways involved in proteostasis, metabolism, and oxidative stress (Fig. 3I, Table S16). The most significant enriched pathways were Bcl2-associated athanogene 2 (BAG2) and Ubiquitin-like protein FAT10 signalling. Complementary STRING analysis identified distinct protein interaction networks (Fig. S2), with clusters of proteins being readily detected associated with chaperone functions, the proteasome, ribosome function, metabolism, sperm morphogenesis and zona pellucida binding. Further DAVID analysis of the core sperm proteome revealed significant enrichment of similar annotation clusters, including chaperone functions, the proteasome, ribosome function, glycolysis, the TCA cycle and flagella. Additional enriched annotation clusters included secretory granules, mitochondrial function and ATP binding. These findings collectively underscore the critical roles of these conserved proteins in ensuring sperm functionality across diverse species.

### Knockout mouse models confirm conserved proteins affect sperm fertilization competency

To investigate the functional relevance of these 135 conserved sperm proteins, we leveraged resources from The International Mouse Phenotyping Consortium (IMPC) database(Dickinson, et al. 2016, Groza, et al. 2022) and the European Mouse Mutant Archive (EMMA)(Hagn, et al. 2007) to obtain knockout (KO) models of these protein-coding genes of interest. This effort yielded five candidates: *Aldh7a1* (Aldehyde dehydrogenase 7 family member A1), *Echs1* (Enoyl-CoA hydratase, short chain 1), *Etfb* (Electron transfer flavoprotein subunit beta), *Ndufa10* (NADH:ubiquinone oxidoreductase subunit A10), and *Pebp4* (Phosphatidylethanolamine binding protein 4). We first established the evolutionary conservation of these proteins across species (Fig. 4A), yielding relatively high conservation between species: Aldh7a1 (median 86.8%), Echs1 (83.4%), Etfb (89.3%), Ndufa10 (77.5%), and Pebp4 (52.3%). Notably species-specific variations were observed for Aldh7a1 (rabbit) and Pebp4 (crocodile).

**Figure 4:**
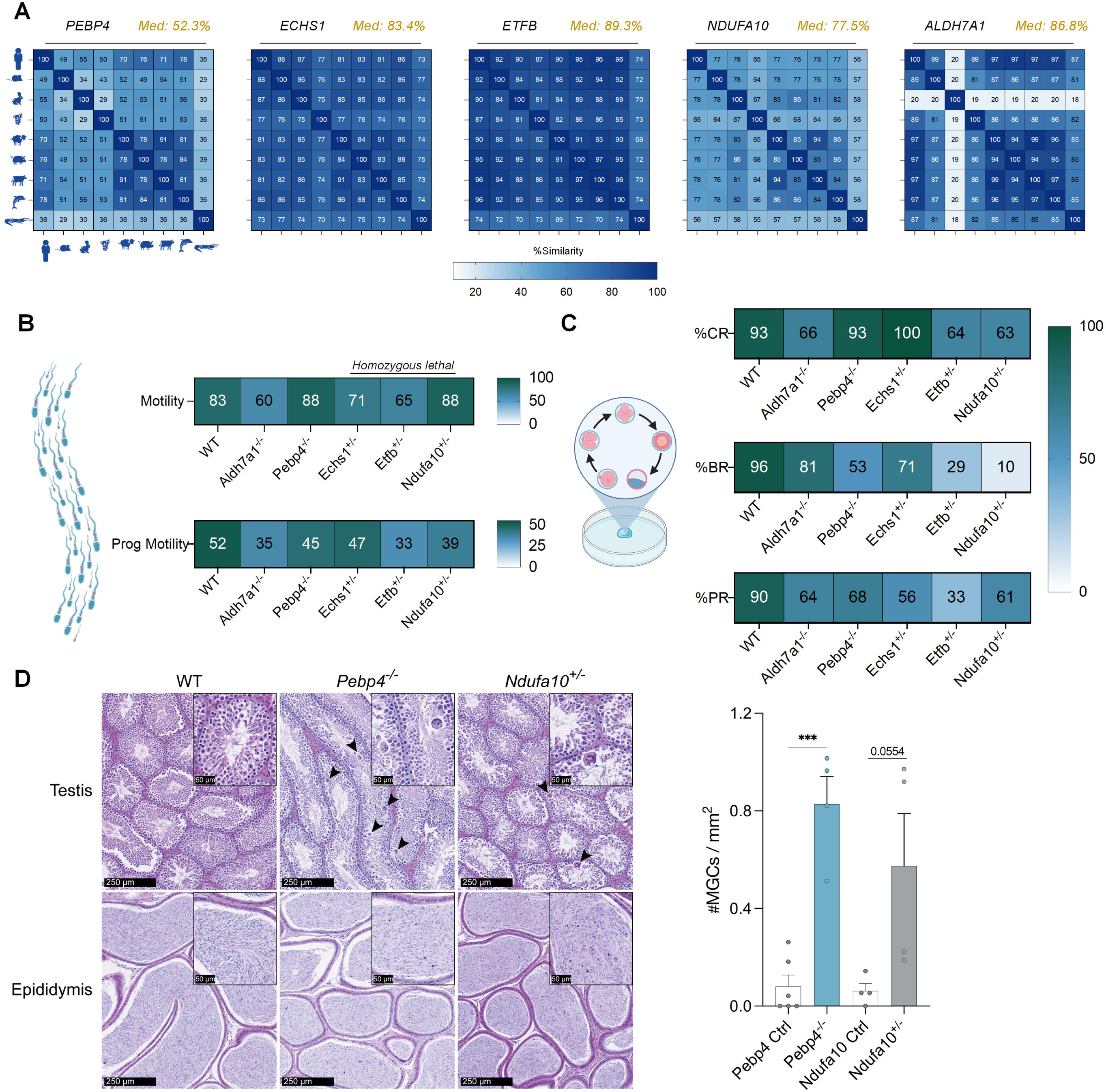
Core sperm protein mice knockouts affect sperm motility and fertilization capacity. (A) UniProt Alignment tool (Clustal Omega) was used to aligned protein sequence for Phosphatidylethanolamine binding protein 4 (Pebp4), Enoyl-CoA hydratase, short chain 1 (Echs1), Electron transfer flavoprotein subunit beta (ETFB), NADH:ubiquinone oxidoreductase subunit A10 (NDUFA10) and Aldehyde dehydrogenase 7 family member A1 (ALDH7A1). Gene knockout (KO) models were generated and sperm from heterozygous males were used for *in-vitro* fertilization (IVF). From these IVF experiments, the heatmaps depict the percentage of (B) motile sperm and those with progressive motility for each KO compared to wildtype (WT). Fertilization capacity was tracked and heatmaps depict the (C) cleavage rate (%CR), blastocyte rate (%BR) and pregnancy rate (%PR). (D) Representative H&E images of the testis and epididymis from WT, *Pebp4*^-/-^, and *Ndufa10*^+/-^ 16-week-old mice, where multinucleated giant cells (MGCs) are indicated by arrowheads. Scale bars = 250 µm and 50 µm for insert image. Quantification of number of MGCs / mm^2^ is represented by a bar chart as Mean ± SEM, with individual datapoints plotted; *** *p* < 0.0001

Through EMMA, we obtained unpublished *in-vitro* fertilization (IVF) data pertaining to heterozygous and homozygous KO mice (Fig. 4). Notably, *Echs1*, *Etfb* and *Ndufa10* are homozygous lethal, and as such, data presented for these genes are from heterozygous males. Total sperm motility analysis of male KO mice showed marked reductions for *Aldh7a1* (23% decrease) and *Etfb* (18%), compared to wildtype (Fig. 4B). Forward progressive motility was further impaired, with proportional reductions of 37% (*Etfb*), 33% (*Aldh7a1*), and 25% (*Ndufa10*) (Fig. 4B). Milder decreases were observed for *Pebp4* (13% loss) and *Echs1* (10% loss). Next, using sperm from these KO males, we performed IVF to evaluate their fertilization potential. The first point of examination was the cleavage rate to the 2-cell stage (Fig. 4C), which mirrored motility trends with marked reductions: *Aldh7a1* (27%), *Etfb* (29%), and *Ndufa10* (30%). Rates for *Pebp4* and *Echs1* remained within expected values. Blastocyst formation rates revealed severe effects, particularly for KOs of *Ndufa10* and *Etfb* with success rates of 10% and 29% respectively (Fig. 4C). Almost halving of success was observed for *Pebp4* (53%), whilst *Aldh7a1* (15%) and *Echs1* (25%) were the least impacted. Pregnancy rates were low for all KOs compared to the expected rate of WTs, with *Etfb* being the most greatly affected (33%), followed by *Echs1* (56%) and remaining KOs ranging from 61% to 68%. Where pregnancy was achieved, litter sizes were within range of expected of age matched wildtype controls (Fig. S3A) with no significant shifts in foetal sex distribution (Fig. S3B).

In complement to the IVF studies, the histology of the male reproductive tract was examined to provide a detailed understanding of its structure and function at microscopic level, and explore any potential contributing effects played by these pivotal tissues. For each KO mouse line, hematoxylin and eosin (H&E) staining was carried out on sections of the testis and epididymis (spermatogenesis (Hermo, et al. 2010a, b) & sperm maturation (Nixon, et al. 2020)), as well as the prostate gland and seminal vesicles (major contributors to the seminal plasma (Robert and Gagnon 1994, Schjenken, et al. 2018)) at 16 weeks of age (Fig. 4D, S4). The *Pebp4* KO displayed the most pronounced abnormalities, including significant presence of multinucleated giant cells (MGC), (symplasts), associated with mild multifocal seminiferous tubular degeneration (Creasy, et al. 2012), and presence of sloughed germ cells in the epididymal caudal tubules, a good indicator of spermatogenic disruption in the testis (De Grava Kempinas and Klinefelter 2014) (Fig. 4D, S4). A similar phenotype of higher number of MGCs was found in *Ndufa10* KOs (Fig.s 4D, S4), but did not achieve significant significance (*p*-value = 0.054), and was accompanied by mild focal vacuolation. Across the five KOs, there were no histopathological changes observed for the prostate or seminal vesicles (Fig. S4), with the exception of *Ndufa10* with hyperplasia in the anterior prostate (or coagulating gland) noted (Fig. S4).

### Conserved proteins loss of function variants linked to human sperm defects

Seeking to further support the clinical relevance of these core sperm proteins, we interrogated our access to nearly 2,900 exomes and genomes of men with quantitative and/or qualitative sperm defects from the MERGE cohort (Stallmeyer, et al. 2024). This analysis returned two men with homo- or hemizygous loss of function (LoF) variants: subject M2218 is homozygous for a stop-gain variant in parkin coregulated: *PACRG*; NM_152410.2:c.369T>A p.(Tyr123Ter). He repeatedly had normal sperm counts but 99-100% sperm head defects and significantly impaired motility. Subject M3692 is homozygous for a frameshift variant in dynein axonemal light intermediate chain 1: *DNALI1*; NM_003462.5:c.490dup p.(Tyr164LeufsTer20). He repeatedly had normal sperm counts but almost all sperm were immotile.

### Characterization of sperm proteins solely identified in different species

The total number of protein IDs for each species was strongly correlated to the number of “unique” (detectable) proteins in that species’ proteome (*r*_s_ = 0.88, *p* = 0.002). While a considerable proportion of proteins in the human (66.3%) and mouse (30.1%) sperm proteomes were only identified in these species, all other species (with much more poorly characterized proteomes) contained <6% proteins not identified in any other species (Table 2, Table S17).

DAVID analysis of unique-to-species proteins highlighted significantly enriched annotation clusters in some species (Table S17). In mice, a range of enriched clusters were observed, including those involved in transcription (RNA splicing (enrichment score (ES) 18.8), small non-coding RNA processing (ES 5.2), RNA helicases (ES 4.2)) and translation (ribosome/mitochondrial translation (ES 8.8). In both cattle and the koala, there was enrichment for secreted proteins (ES 4.3, 3.5 respectively), however the proteins within these clusters were species-specific. Secreted proteins only identified in the cattle sperm proteome included beta defensins (DEFB108B, DEFB116), cytokines (IL12B, IL34, CCL2, CTF1), proteins with antimicrobial activity (VIP, ADM) and RNase 1. In contrast, the enriched cluster of secreted proteins only identified in the koala sperm proteome largely contained pro-hormones (AVP, RLN2, PTH, INSL5, POMC). Proteins uniquely identified in dolphin sperm showed weak but significant enrichment of functions associated with cell and gonadal differentiation (ES 2.6), including AMH, TSPY2 and TSPY8. There were no significantly enriched annotation clusters in unique-to-species proteins in the boar, rabbit, sheep or crocodile sperm proteomes.

As described above, depending on the species, 0.9 – 4.1% of protein identifications were not successfully converted to humanized IDs for further analysis. While many of these were poorly characterized proteins that may indeed have human equivalents, it is likely that some of these represent additional unique-to-species proteins. Examples of such proteins that are thought to be specific to one or several closely related species (albeit most with homologues in more diverse species) include mouse seminal vesicle proteins (SVS3A, SVS3B, SVS4, SVS5, SVS6)(Karn, et al. 2008), boar carbohydrate-binding protein AQN-1(Kraus, et al. 2005), bovine spermadhesin Z13 (Haase, et al. 2005), and rabbit semen coagulum protein (SVP200) (Lundwall, et al. 2020).

### Reproductive strategies reflected in sperm proteomes

In seeking to investigate the influence of evolutionary and imposed reproductive strategies on sperm protein composition, we further interrogated each species proteome by focusing on sperm metabolism preference (glycolysis preference vs no preference), location of the testes (internal vs external), and history of selective breeding (yes vs no). Species were stratified into their appropriate groups and Venn Diagram analyses were conducted to determine the proteins unique to each biological context. Firstly, focusing on sperm metabolism, a comparison of species with a preference for glycolytic sperm metabolism against those with no preference between glycolysis and oxidative phosphorylation, resulted in a 49.7% shared overlap of proteins (1,857) (Fig. 5A). The larger component of this comparison was those species with no sperm metabolism preference (cattle, rabbit, sheep), with 1,385 unique proteins. Only species that preferentially use glycolysis showed enrichment for a variety of metabolically relevant pathways, including degradation of amino acids (valine, isoleucine, tryptophan, leucine, cysteine, phenylalanine, alanine, glutamine) and N-acetylglucosamine (Table S18).

**Figure 5:**
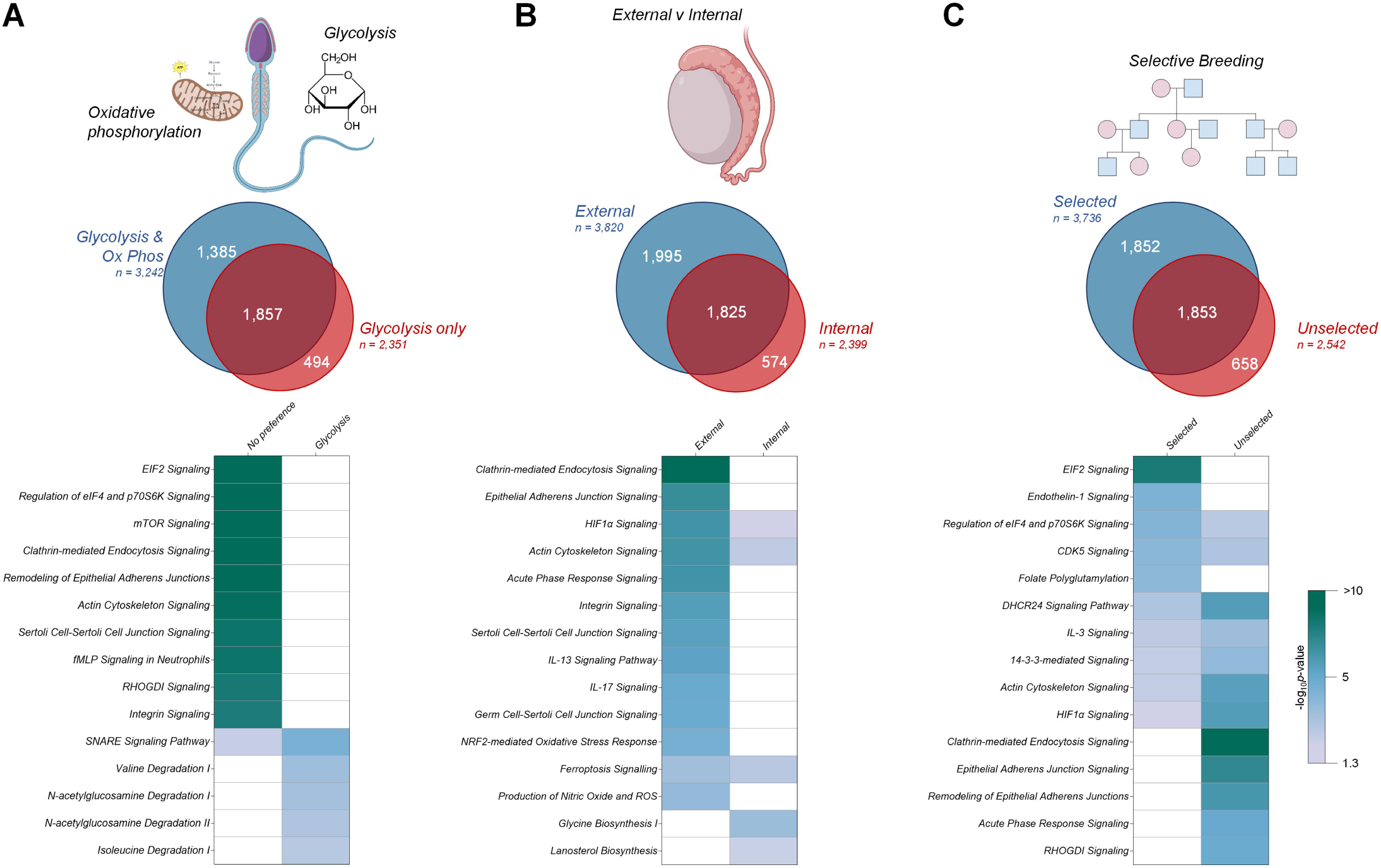
Reproductive strategies analyses. Sperm proteomes were stratified into three analyses, investigating the influence of evolutionary and imposed reproductive strategies on sperm protein composition, focusing on (A) sperm metabolism preference (glycolysis preference vs no preference), (B) location of the testes (internal vs external), and (C) history of selective breeding (yes vs no). Each analyses included Venn Diagrams to determine unique proteins to each biological context, which were further subject to analysis using Ingenuity Pathway Analysis (IPA). Heatmaps depict the comparative analysis of the resultant molecular functions.

A subsequent comparison of sperm proteomes based on species with internal vs external testes revealed 41.5% of proteins (1,825) were shared, with only 13.1% (574) proteins unique to species with internal testes (crocodile, dolphin) (Fig. 5B). Species with external testes showed significant enrichment for signalling involving a range of interleukins (e.g. IL-3, −13, −17), reactive oxygen species production and NRF2-mediated oxidative stress response pathways, as well as clathrin mediated endocytosis signalling, adherens junction signalling, HIF1α signalling, acute phase response signalling, and integrin signalling. By comparison, species with internal testes showed significant enrichment for pathways involved in glycine and cholesterol synthesis (Table S18).

Lastly, species were compared based on whether they have a history of selective breeding; a strategy that revealed 42.9% (1,853) proteins were shared regardless of breeding practices (Fig. 5C). However, species under selective breeding pressures (cattle, boar, rabbit, sheep) yielded the greater number of unique proteins amounting to a total of 1,853 proteins (42.1%). Conversely, sperm from those species without selective pressures were associated with the unique expression of 658 proteins (8.2%). Selectively bred species showed higher enrichment for endothelin-1, EIF2 and CDK5 signalling. In comparison, species with no history of selective breeding showed higher enrichment for adherens junction signalling (Table S18). Unique and overlapping protein IDs for each of the 3 biological contexts comparisons are supplied in Table S18.

## DISCUSSION

From a total of 29 datasets across 12 vertebrate species (>2TB of RAW data), we have generated the most comprehensive cross species analysis of the mature sperm proteome to date, identifying a grand total of 13,853 unique proteins residing in the sperm of those species studied herein. Within this dataset, we discerned a core sperm proteome of 45 proteins conserved at the species level and 135 proteins conserved at the order level, underscoring a fundamental molecular framework essential for generating fertilization-competent spermatozoa. Enrichment analyses linked these core proteins to critical pathways in proteostasis and sperm metabolism, while knockout mouse models of selected conserved proteins confirmed their influential roles in governing sperm motility, fertilization success, and overall reproductive health. Together, our findings demonstrate that despite the remarkable diversity in reproductive strategies observed across vertebrates, including differences in sperm metabolic preferences, testicular location, and histories of selective breeding, a fundamental set of molecular components remain steadfastly conserved. This universal baseline provides a platform upon which species-specific adaptations are layered, shaping the unique sperm proteomes that ultimately define each organism’s reproductive biology.

The uptake of proteomics technologies in reproductive biology is ever expanding, with over 20-fold increase in 20 years,(Skerrett-Byrne, et al. 2024) as improvements to mass spectrometry (MS) technology and bioinformatic pipelines continue to explode. As a field, proteomics is reaching an intriguing point, as these RAW datasets can be viewed as ‘*living datasets*’, from which we can continuously yield new insights as bioinformatic tools, database annotations, and computational capacities advance.(Dai, et al. 2024, Drew, et al. 2017) Whilst there are several variables affecting their ‘*mine-ability*’, early analyses often only scratch the surface of what these complex spectra contain; proteins may remain unidentified simply because the necessary reference sequences or annotation frameworks are inadequately annotated at the time.(Willems, et al. 2020) With decreasing costs of whole-genome sequencing, genome assemblies are improving, particularly across diverse taxa. This, coupled to ever evolving algorithms for spectral matching and integrative proteomics workflows, is beginning to reveal novel proteins, post-translational modifications, or subtle quantitative differences that were once hidden.(Polasky, et al. 2023, Vaudel, et al. 2016, Willems, et al. 2020) Thus, the continuous reanalysis of MS datasets is not merely a luxury, but a crucial endeavour to fully leverage these rich repositories of biological information, ensuring that the scientific community continues to extract meaningful knowledge from the collective body of proteomic data. In complement to this, large-scale proteomic atlases, such as the recently expanded draft of the human and mouse proteomes,(Giansanti, et al. 2022, Wang, et al. 2019) providing a systematic framework for characterising protein landscapes across species. Adapting such approaches to non-traditional species, alongside improving genome sequencing efforts, will be essential to fully capture the complexity of sperm proteomes across diverse taxa.

Of the >2,000 sperm proteome studies found on PubMed,(Skerrett-Byrne, et al. 2024) a mere 218 studies have data deposited in a publicly available repository, a poor return of ∼10%. To our knowledge there have yet to be any studies reanalysing the RAW MS data of previous studies focused on mature sperm cells. Leveraging this unique situation and technological advancements, here, we have demonstrated the capacity to assemble proteomic profiles for 12 taxa, achieving an unprecedented depth of coverage and unveiling numerous unique and conserved proteins. Comparing the proteomic depths of the studies included to these new sperm proteomes, we demonstrate improved depths from subtle increases in rabbit (initially 1,360 proteins(Juárez, et al. 2020), now 1,605; 18% increase) to more overt increases in the crocodile (initially 1,119 proteins(Nixon, et al. 2019), now 2,855; 155% increase). Despite these strides, the proteomic annotation of non-model species still lags behind that of humans and established model organisms, where genome annotation and protein databases are significantly more mature. This is highlighted by the discrepancy between the human and boar sperm proteome, the latter of which has only 35% the proteome size of the former, and even more dramatically with the koala being only 6.3% the size of the human sperm proteome. Consequently, the wealth of newly identified proteins reported here, while indicative of greater analytical depth, is still constrained by the uneven quality of reference data, limiting our ability to conclusively determine whether certain proteins are genuinely unique to a given species or simply undetected in others due to inadequate sampling or annotation. This highlights the need for greater annotation placed upon non-model species in NCBI and UniProt databases, and the pursuit of new analytical tools.(Heck and Neely 2020, Van den Broeck, et al. 2023)

Interestingly, even beyond direct protein-to-protein comparisons, our functional analyses highlight complex patterns of convergence and divergence that transcend traditional phylogenetic boundaries. For instance, despite their substantial evolutionary distance, koalas and sheep emerge as functionally aligned in terms of pathways underpinning their sperm proteomes, underscoring the powerful influence of reproductive strategies and physiological demands on proteome composition. Conversely, we observed species-specific pathways enrichment, with crocodiles displaying the great enrichment of fatty acid β-oxidation, an important source of long-term sustained energy as sperm ascends several meters of the female reproductive tract (Gist, et al. 2008, Nixon, et al. 2019). These findings not only illuminate previously unknown dimensions of sperm biology but also point toward an exciting future in which comparative proteomics will continue to uncover unexpected connections and functional adaptations, broadening our understanding of reproductive strategies and their molecular underpinnings.

The identification of a core sperm proteome comprising 45 proteins at the species level and 135 at the order level underscores the presence of a conserved molecular foundation essential for sperm function across evolutionary lineages. Notably, given the size of the smallest proteome, the koala (556 proteins), these conserved proteins represent a 24.3% level of conservation at the order level, highlighting the functional importance of these proteins despite proteome size variation. Focusing on the sequence conservation of these core sperm proteins, we observed a median of 94.4% conservation at the amino acid level. Proteins such as heat shock protein A2 (HSPA2) and voltage dependent anion channel 3 (VDAC3) maintain exceptionally high sequence conservation, reflecting their critical and broadly indispensable roles in processes like protein folding, sperm maturation, and capacitation.(Arcelay, et al. 2008, Li, et al. 2023, Nixon, et al. 2017, Redgrove, et al. 2012, Sampson, et al. 2001) By contrast, proteins critical for zona pellucida binding and penetration, acrosin (ACR)(Dudkiewicz 1983, Hua, et al. 2023, Liu and Gordon Baker 1993), zona pellucida-binding protein (ZPBP),(Dun, et al. 2010, Lin, et al. 2007) and calcium-binding tyrosine-phosphorylation regulated protein (CABYR),(Naaby-Hansen, et al. 2002, Skerrett-Byrne, et al. 2022) exhibit greater sequence variability, perhaps signifying adaptations to specific reproductive environments or selective pressures.

This is supported by previous work which sought to compare the sperm proteome within three closely related *Mus* species that experience different level of sexual selection(Vicens, et al. 2017). Whilst not a defined difference in protein sequence, this work highlighted significant interspecific protein abundance divergence of proteins which govern sperm–egg interactions, including ACR, ZPBPs, and conversely no significant shifts in HSPA2 or protein kinase cAMP-dependent type I regulatory subunit alpha (PRKAR1A). Seeking to understand how these core sperm proteins may interact, *in-silico* enrichment analyses grouped them into known networks integral to sperm motility (flagellar motility), energy source of the sperm cell (oxidative phosphorylation [OXPHOS]), and fertilization, encompassing zona pellucida binding and acrosome reaction. A key study from 2016, included in our work, carried out proteomics on sperm collected from 19 placental mammalian species, identified a core sperm proteome of 623 proteins (Bayram, et al. 2016). Importantly, this study leverages two advantages: 1) comparatively narrower range of species, ungulates and rodents; 2) all proteomics sample preparation, MS analysis and data processing are uniform. However, there is considerable overlap with our study in regards to the significantly enriched pathways and functions which connect the core sperm proteome, namely metabolic processes such as OXPHOS, glycolysis and the tricarboxylic acid cycle (TCA), acrosome assembly, zone pellucida binding and proteasome function (Bayram, et al. 2016). Notably, the emergence of the highly enriched proteostasis-associated pathways, involving proteins like BAG2 and FAT10, provides intriguing hints of previously underappreciated quality control processes that may safeguard sperm function by ensuring proper protein folding, timely degradation, regulation of capacitation and acrosomal reaction, thereby facilitating successful fertilization.(Cafe, et al. 2021, Smyth, et al. 2024) For instance, the enrichment of BAG2 in the conserved sperm proteome, known to modulate HSPA2 in spermatogenesis,(Yin, et al. 2020) raises the potential of a post-testicular role in maintaining HSPA2 functionality in facilitating sperm-egg adhesion and binding.(Nixon, et al. 2015, Smyth, et al. 2024)

Seeking to build upon this important database (https://reproproteomics.shinyapps.io/ShinySpermKingdom/) and demonstrate how these analyses can help build new knowledge on sperm biology, we were granted special access to the European Mouse Mutant Archive (EMMA)(Hagn, et al. 2007) to obtain sperm and *in vitro* fertilization (IVF) data, from knockout (KO) mouse models targeting selected proteins from the core proteome. We elected to focus on proteins which had not previously been implicated in sperm morphology or functional maturation; metabolic enzyme Aldh7a1 (Aldehyde dehydrogenase 7 family member A1)(Korasick and Tanner 2021); mitochondrial enzymes Echs1 (Enoyl-CoA hydratase, short chain 1)(Burgin and McKenzie 2020) and Etfb (Electron transfer flavoprotein subunit beta)(Henriques, et al. 2021); mitochondrial complex I subunit Ndufa10 (NADH:ubiquinone oxidoreductase subunit A10) (Formosa, et al. 2018); and a member of the phosphatidylethanolamine-binding proteins, Pebp4, although detected in bull sperm and semen previously,(An, et al. 2012, Somashekar, et al. 2017) no studies have demonstrated its role in sperm function. Here for the first time, KOs of these five core genes have been shown to directly influence sperm quality, motility, and fertilization potential. Intriguingly, the largest decreases observed in total sperm motility and progressive motility were associated with the four proteins located in the mitochondrial matrix, each of which play key roles in mitochondrial function. Given the pivotal role mitochondria play in the energy required to drive sperm motility(Piomboni, et al. 2012), it lends to reason these proteins may have regulatory roles in ensuring efficient OXPHOS/ATP production in sperm mitochondria. Further to the KO mouse work, we sought to investigate the clinical relevance of these proteins in the context of human infertility. Utilising the MERGE cohort, we queried our 135 conserved sperm proteins against the almost 2,900 males with quantitative and/or qualitative sperm defects, identifying two men with homozygous loss-of-function variants in *PACRG* and *DNALI1*. Notably, a recent mouse KO study demonstrated that PACRG interacts within a manchette-associated complex, and is essential for proper sperm assembly (Yap, et al. 2023). Moreover, KO mice exhibited a significant reduction in sperm count, with the remaining sperm characterized by abnormally shaped heads and bent tails. DNALI1 is a component of the inner dynein arms and in men affected by biallelic pathogenic variants, immotile sperm exhibiting an asymmetric fibrous sheath of the flagella have been described (Sha, et al. 2022). Together, these experimental models lend strong empirical support to the notion that the core sperm proteome is not merely a historical residue of evolutionary conservation, but rather a functional blueprint vital for the generation and maintenance of fertilization-competent sperm across species. Further studies are underway to tease apart the contributions of the remaining 130 conserved proteins and whether there are compensatory mechanisms in play.

Beyond their universally conserved core, sperm proteomes also reflect diverse biological contexts that shape reproductive strategies and outcomes. For instance, contrasting testicular environments, as represented by species with either external or internal testes, impart distinct proteomic signatures on mature sperm. Species with external testes exhibit enrichments in signalling pathways associated with interleukin-mediated communication, reactive oxygen species management, and NRF2-mediated oxidative stress responses, as well as pathways related to cell-cell interaction and endocytic processes. These findings suggest that externalized gonads, potentially evolving under selective pressures linked to temperature regulation or hypoxic conditions, have prompted the refinement of sperm’s molecular toolkit to bolster resilience and maintain fertilisation capacity. Conversely, species with internal testes show a bias toward pathways involving glycine and cholesterol synthesis, hinting at metabolic adjustments that support sperm function in a more thermally stable yet potentially resource-limited environment.

Similarly, historical selective breeding regimes and species-specific metabolic preferences leave distinguishable marks on the sperm proteome. The enrichment of pathways associated with microRNA biogenesis and endothelin-1 signalling in selectively bred species indicates that human-driven selection can influence the molecular composition of sperm, potentially altering traits related to sperm maturation, capacitation, or early embryonic development through epigenetic and signalling mechanisms.(Conine and Rando 2022, Sharma, et al. 2018, Spadafora 2023, Tomar, et al. 2024, Trigg, et al. 2021) In contrast, non-selected species’ enrichment for adherens junction signalling points to a more basal state of sperm cell-cell interaction and membrane integrity.(Lui, et al. 2003, Wen, et al. 2016) Additionally, the preferential use of glycolysis as an energy source(du Plessis, et al. 2015) in certain taxa is mirrored by an increased representation of amino acid degradation and N-acetylglucosamine metabolism pathways. Such metabolic footprints underscore how ecological and evolutionary pressures, ranging from the physical layout of the male reproductive tract to dietary constraints and energy demands, coalesce to shape the molecular identity of sperm across species. In essence, the sperm proteome reflects not only a shared, evolutionarily ancient blueprint but also the distinct biological contexts that steer reproductive strategies and outcomes in varied environmental and evolutionary landscapes. Complementing the full breadth of analyses discussed here, we have deployed a Shiny application, ShinySpermKingdom (https://reproproteomics.shinyapps.io/ShinySpermKingdom/), to support the accessibility and interpretability of these datasets beyond static documents, allowing for effective data-driven insights by researchers in the field (Skerrett-Byrne, et al. 2024).

Despite the breadth and depth of the current dataset, several limitations constrain the interpretation and generalisability of our findings. Foremost, the use of publicly available proteomic data, generated over a span of years and employing diverse sample preparation protocols, mass spectrometry platforms, and analytical pipelines, precludes direct, quantitative ‘*apples to apples*’ comparisons between species. Although advancements in bioinformatics and the curation of comprehensive repositories have greatly expanded the taxonomic breadth of sperm proteome studies, many datasets remain hampered by incomplete metadata, inconsistent reporting standards, and limited raw data availability. These constraints hinder reproducibility, limit standardized reanalysis, and reduce the clarity of species-to-species comparisons. Additionally, our reliance on human orthologues for pathway analyses, while essential for leveraging robust functional annotation tools, inevitably restricts our ability to fully explore lineage-specific adaptations or signatures of selective pressure acting on particular sperm proteins. Similarly, we have focused exclusively on chordate species, thus excluding the vast diversity of reproductive strategies found in invertebrates and other non-chordate taxa. Additionally, the marked disparity in proteome sizes across species highlights that coverage bias remains a significant concern. This discrepancy, exemplified by the koala sperm proteome relative to that of humans or mice, likely reflects the limits of detection, underscored by poor annotation of these non-model species. As proteomic technologies and genome annotations continue to improve for these underrepresented species, many more proteins will inevitably be discovered, reinforcing the argument that our current dataset offers a glimpse of the full sperm proteome complexity. Recognizing these acquisition biases paves the way for new funding proposals aimed at refining reference annotations and expanding high-quality MS coverage across diverse species. By systematically addressing these limitations, the field can move closer to an accurate and comprehensive understanding of sperm proteomes at a global scale. Until then, our findings, while important and innovative, should be viewed as a framework upon which to build as the global sperm proteome landscape becomes more thoroughly mapped and understood.

Future investigations will benefit from a more controlled and integrated experimental framework, enabling true comparisons across species. Standardising sample collection protocols, employing uniform mass spectrometry methodologies, and harmonising data processing pipelines will help overcome current disparities in data quality and annotation. As proteome databases are further refined, and as more non-model species achieve higher-quality genome assemblies, future analyses will be better positioned to distinguish genuine species-specific proteins from those simply missing in incomplete datasets. Additionally, exploring the role of post-translational modifications (PTMs), such as phosphorylation, acetylation, and glycosylation, across multiple taxa will be critical for understanding how subtle regulatory mechanisms modulate sperm functionality. Indeed, since sperm protein composition is largely defined during epididymal maturation,(Skerrett-Byrne, et al. 2022) incorporating temporal and spatial sampling strategies alongside PTM profiling may reveal how species adapt their sperm at the molecular level to an array of environmental pressures.

Ultimately, comprehensive multispecies studies employing rigorous and consistent workflows have the potential to create living, expandable proteomic databases that will evolve as analytical capabilities and reference annotations improve. By repeatedly revisiting and reanalysing mass spectrometry datasets, the field can unlock layers of complexity that have, until now, remained concealed. Such iterative proteomic approaches, informed by emerging knowledge of sperm physiology and integrated with genomic, transcriptomic, and epigenomic data, will open new avenues for understanding the evolutionary and functional contexts of sperm proteomes. Here, we demonstrate the power of systematically revisiting and reanalysing proteomic data, identifying a set of 135 conserved proteins critical to sperm function, including several that have never been previously implicated in sperm biology. This study underscores the importance of continuous proteomic refinement, allowing for the identification of previously overlooked but evolutionally and functionally essential proteins. By embracing this approach, future research to not only refine our grasp of the universal underpinnings of sperm biology but also reveal how species-specific adaptations arise and shape reproductive success across diverse taxa.

## AUTHOR CONTRIBUTIONS

Conceptualisation, T.P., B.N., and D.A.S.B.; Methodology, T.P. and D.A.S.B.; Software, D.A.S.B.; Investigation, T.P., B.N., T.K., R.T., A.S.M., P.S.B., F.T., T.S., S.K., V.G.D., H.F., S.M., M.H.A., and D.A.S.B.; Formal Analysis D.A.S.B.; Validation R.T., A.S.M., P.S.B., V.G.D., H.F., S.M., M.H.A., and D.A.S.B; Visualisation, T.P., and D.A.S.B; Writing – Original Draft, T.P., and D.A.S.B; Writing – Review & Editing, B.N., T.K., R.T., A.S.M., P.S.B., F.T., T.S., S.K., V.G.D., H.F., S.M., M.H.A.; Funding Acquisition, B.N. and D.A.S.B; Resources, B.N. R.T., F.T., T.S., S.K., V.G.D., H.F., S.M., M.H.A., and D.A.S.B.; Supervision, B.N. and D.A.S.B.

## ACKNOWLEDGEMENTS

We thank the Academic and Research Computing Support team, The University of Newcastle who provided High Performance Computing Infrastructure to support the bioinformatics analyses. We also thank David MacKenzie for their graphic design input and support. We thank, S. Dunst and B. Rey, for technical support with the sperm and IVF culture experiments. We thank the technicians and animal caretakers of the German Mouse Clinic. We thank Dr Wei Zhou for his insightful questions at The Society for Reproductive Biology Annual Scientific Meeting.

## Supplemental Figures

**Figure S1:**
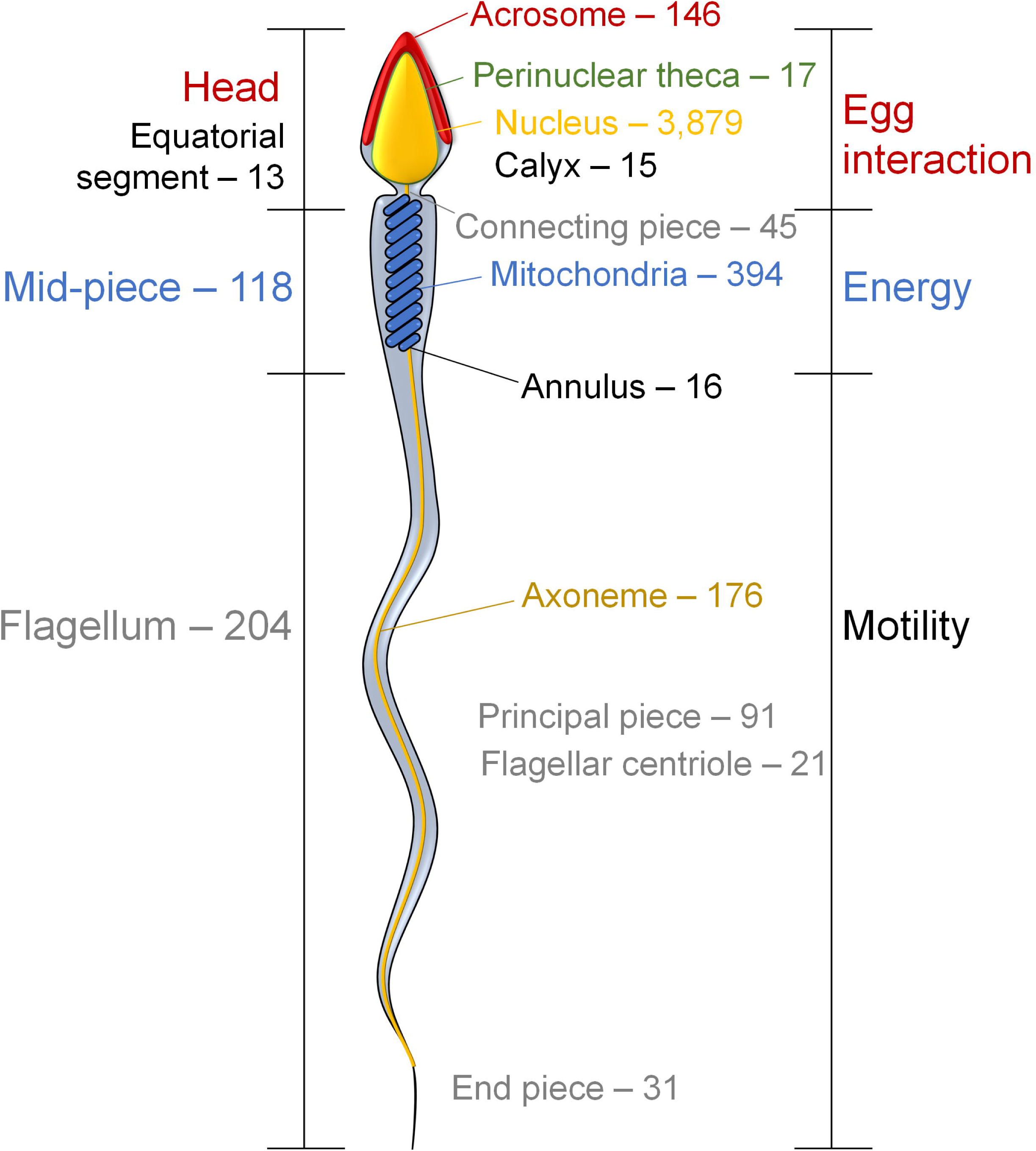
Sperm protein localization. Interrogation with UniProt maps proteins to their known sperm localizations.

**Figure S2:**
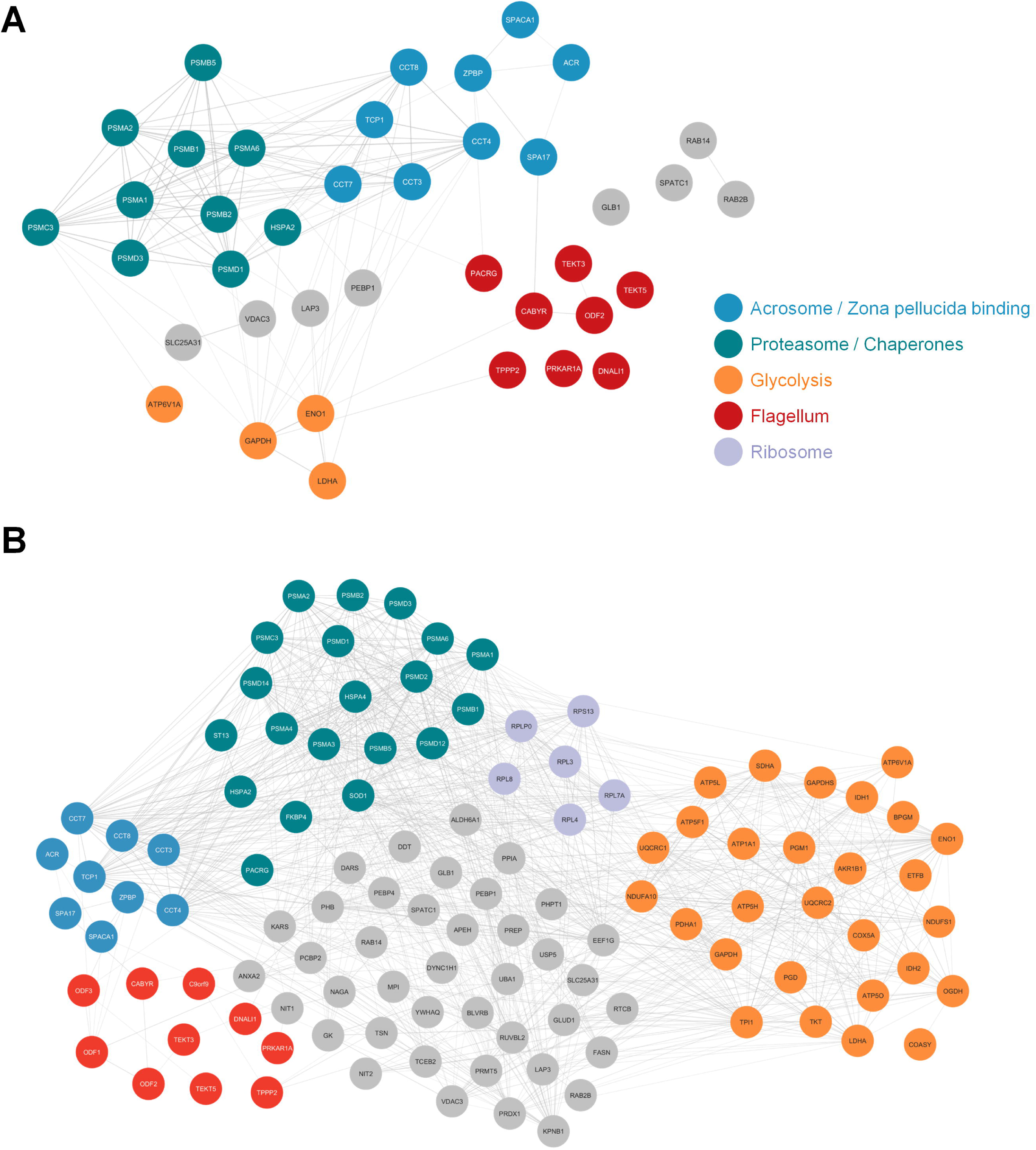
Protein-to-protein interaction networks of core sperm proteomes. STRING network analyses of the core sperm proteome at the (A) species (45 proteins) and (B) order (135 proteins) taxonomic levels, drawn and coloured with Cytoscape.

**Figure S3:**
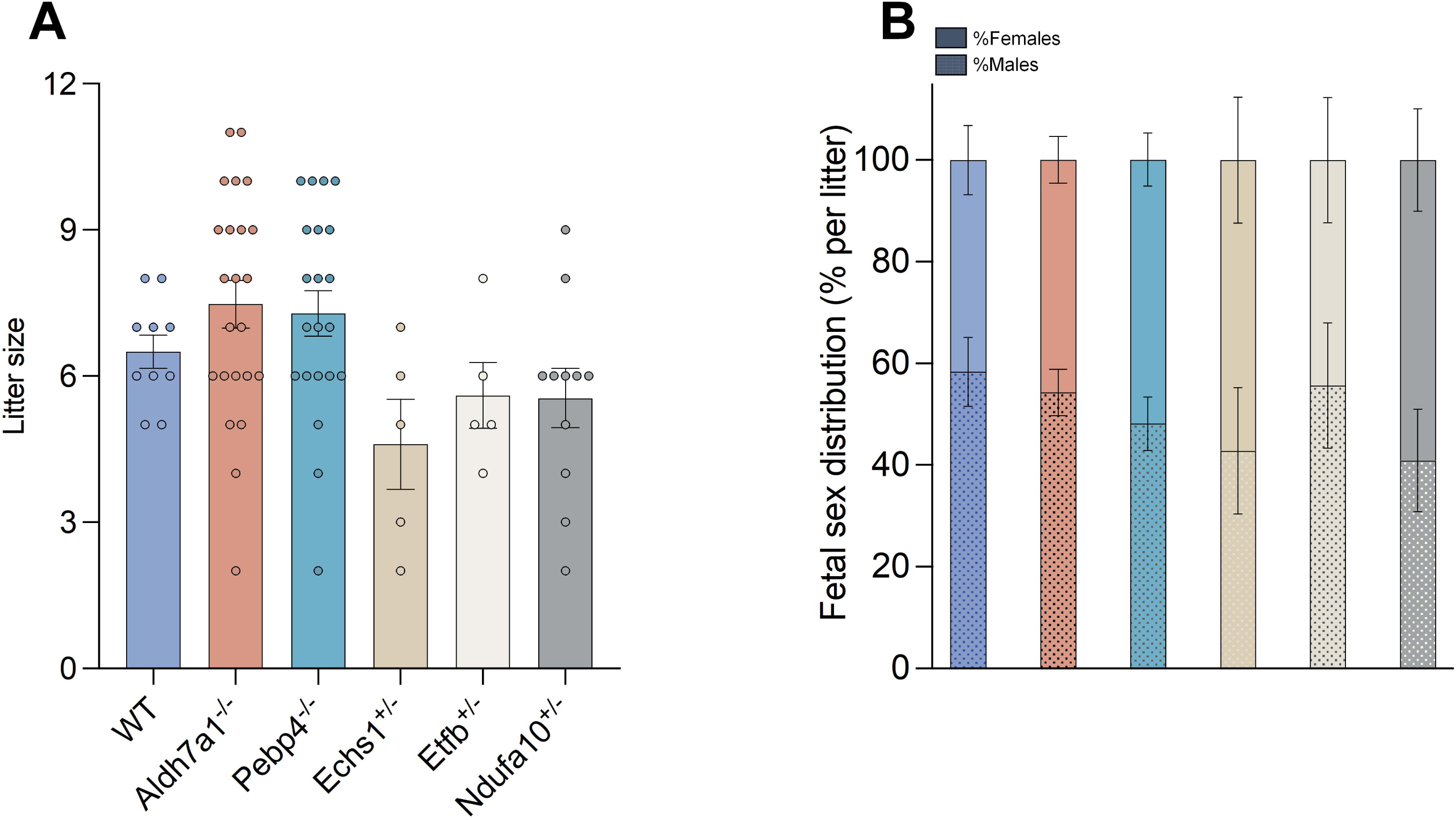
Knockout mouse model pregnancy outcomes. Following successful births, the (A) litter size and (B) foetal sex distribution was recorded.

**Figure S4:**
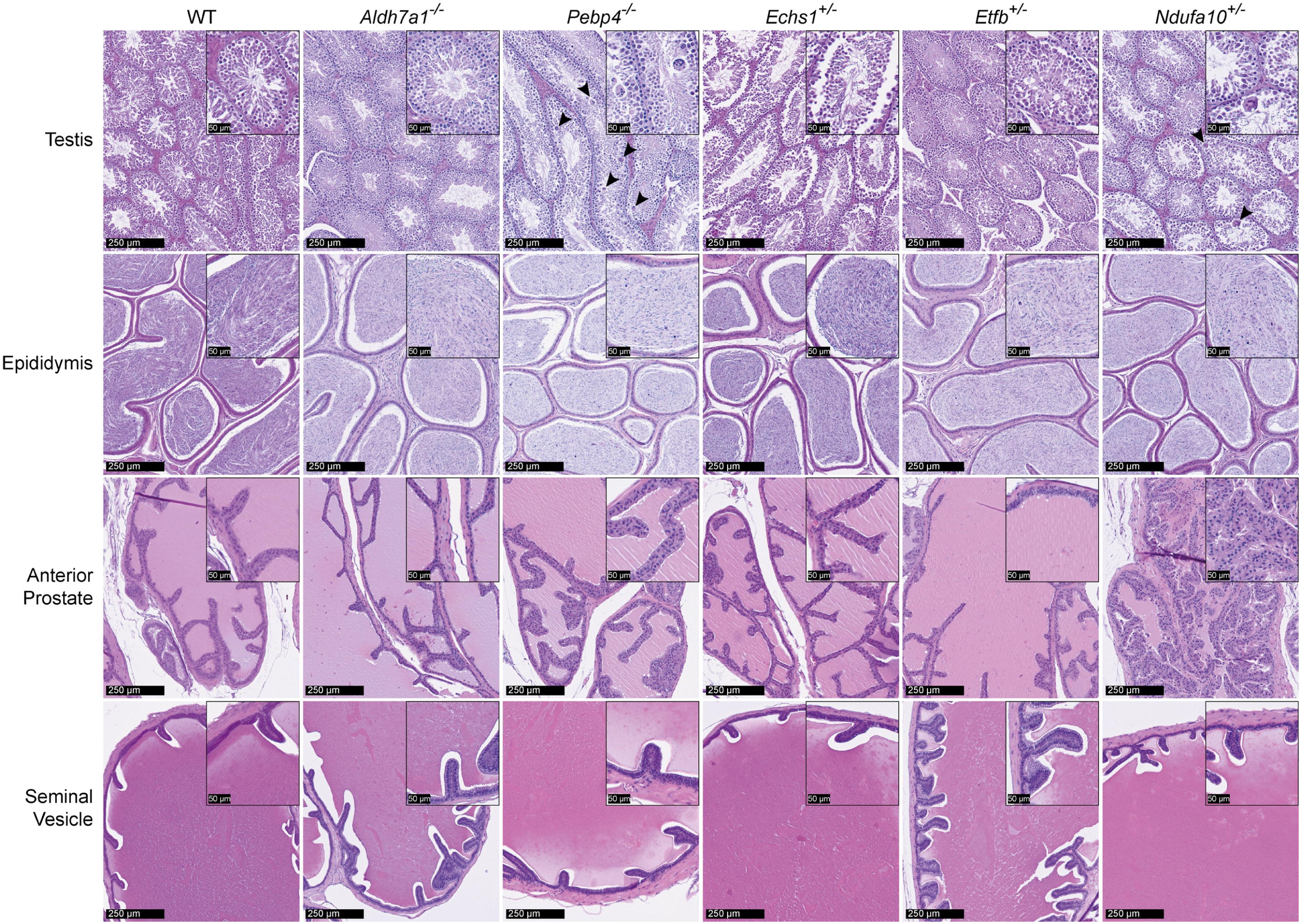
Histopathology of five KO models across the male reproductive tract. Representative images of H&E-stained sections from testis, cauda epididymis, anterior prostate and seminal vesicles of controls and knockout mice for the following genes: Aldehyde dehydrogenase 7 family member A1 (*Aldh7a1*), Phosphatidylethanolamine binding protein 4 (*Pebp4*), Enoyl-CoA hydratase, short chain 1 (*Echs1*), Electron transfer flavoprotein subunit beta (*Etfb*), and NADH:ubiquinone oxidoreductase subunit A10 (*Ndufa10*). Mice were 16 weeks old in all lines except for *Aldh7a1* (19 weeks old). Multinucleated giant cells (MGCs) are indicated by arrowheads. Scale bars are 250 µm and 50 µm.

## Supplemental Tables

**Table S1**

Complete publicly available proteome datasets investigated in this study. Also includes information on the FASTA files used across species.

**Tables S2-13**

The refined proteomic list for all 12 species, including UniProt accession, protein name, gene, level of annotated evidence, reviewed status and function from UniProt.

**Table S14**

All humanized proteomic datasets.

**Table S15**

The 45 proteins conserved at the species level and 135 proteins conserved at the order level.

**Table S16**

Core sperm proteome analysis outputs from Ingenuity Pathway Analysis (IPA)

**Table S17**

All proteins uniquely detected within each species and their respective outputs from Database for Annotation, Visualization and Integrated Discovery (DAVID).

**Table S18**

The analyses pertaining to the biological context comparisons, with unique proteins identifications listed, alongside the functional outputs from Ingenuity Pathway Analysis (IPA).

